# Cardiomyocyte BRAF and type 1 RAF inhibitors promote cardiomyocyte and cardiac hypertrophy in mice *in vivo*

**DOI:** 10.1101/2021.08.14.455637

**Authors:** A Clerk, DN Meijles, MA Hardyman, SJ Fuller, SP Chothani, JJ Cull, STE Cooper, HO Alharbi, K Vanezis, LE Felkin, T Markou, SJ Leonard, SW Shaw, OJL Rackham, SA Cook, PE Glennon, MN Sheppard, JC Sembrat, M Rojas, CF McTiernan, PJ Barton, PH Sugden

## Abstract

The extracellular signal-regulated kinase 1/2 (ERK1/2) cascade promotes cardiomyocyte hypertrophy and is cardioprotective, with the three RAF kinases forming a node for signal integration. Our aims were to determine if BRAF is relevant for human heart failure, if BRAF promotes cardiomyocyte hypertrophy, and if Type 1 RAF inhibitors developed for cancer (that paradoxically activate ERK1/2 at low concentrations: the “RAF paradox”) may have the same effect. BRAF was upregulated in heart samples from patients with heart failure compared with normal controls. We assessed the effects of activated BRAF in the heart using mice with tamoxifen-activated Cre for cardiomyocyte-specific knock-in of the activating V600E mutation into the endogenous gene. We used echocardiography to measure cardiac dimensions/function. Cardiomyocyte BRAF^V600E^ induced cardiac hypertrophy within 10 d, resulting in increased ejection fraction and fractional shortening over 6 weeks. This was associated with increased cardiomyocyte size without significant fibrosis, consistent with compensated hypertrophy. The experimental Type 1 RAF inhibitor, SB590885, and/or encorafenib (a RAF inhibitor used clinically) increased ERK1/2 phosphorylation in cardiomyocytes, and promoted hypertrophy, consistent with a “RAF paradox” effect. Both promoted cardiac hypertrophy in mouse hearts *in vivo*, with increased cardiomyocyte size and no overt fibrosis. In conclusion, BRAF potentially plays an important role in human failing hearts, activation of BRAF is sufficient to induce hypertrophy, and Type 1 RAF inhibitors promote hypertrophy via the “RAF paradox”. Cardiac hypertrophy resulting from these interventions was not associated with pathological features, suggesting that Type 1 RAF inhibitors may be useful to boost cardiomyocyte function.

## Introduction

Mammalian cardiomyocytes (the contractile cells of the heart) withdraw from the cell cycle in the perinatal period and become terminally-differentiated [1]. However, the adult heart is subjected to pathophysiological stresses that increase workload and require greater cardiac output. To accommodate this, cardiomyocytes undergo hypertrophic growth (increasing in size in the absence of proliferation), increasing and adapting the myofibrillar apparatus, leading to cardiac hypertrophy. In some instances, this is a benign adaptation to, for example, pregnancy or endurance exercise [2]. Such “physiological” hypertrophy is generally reversible once the stress has ceased, and is not associated with fibrosis. In chronic conditions (e.g. hypertension), or with acute damage to the heart (e.g. a result of myocardial infarction), the heart initially adapts to maintain cardiac output with cardiomyocyte hypertrophy, but this adaptive hypertrophy is not sustained. Contractile function becomes compromised and heart failure develops, with increased cardiomyocyte cell death [3], loss of capillaries [4], inflammation and fibrosis [5].

The extracellular signal-regulated kinase 1/2 (ERK1/2) cascade is the prototypic mitogen-activated protein kinase (MAPK) pathway, first identified in the context of its role in promoting cell division in proliferating cells. However, this pathway also promotes cardiomyocyte hypertrophy and is cardioprotective [6-8]. ERK1/2 operate in a cascade in which RAF kinases (ARAF, BRAF, RAF1) phosphorylate and activate MAPK kinases 1/2 (MKK1/2), and these phosphorylate/activate ERK1/2 [9]. RAF kinases receive inputs from different stimuli and are regulated at multiple levels [9]. Activation requires interaction with activated Ras and, in proliferating cells, this promotes formation of active RAF homo- or heterodimers. RAF kinases are subject to both activating and inhibitory phosphorylations. In particular, RAF1 requires phosphorylation of Ser338/Tyr341 for activity (all phosphorylation sites refer to the human protein; equivalent residues are present in the mouse and rat proteins). BRAF has high basal activity because it possesses acidic residues in this domain, but is phosphorylated on Ser445 [equivalent to RAF1(Ser338)].

The regulation of RAF kinases in terminally-differentiated cardiomyocytes is less well understood. RAF1 and ARAF are activated in cardiomyocytes by growth factors and hypertrophic agonists such as endothelin-1 [10]. As in proliferating cells, RAF1 activation requires activation of the small G protein, Ras, and this leads to phosphorylation of RAF1(Ser338) [11, 12]. Cardiomyocyte-specific expression of dominant-negative RAF1 in mice increases cardiomyocyte apoptosis, with cardiomyopathy developing in response to pressure-overload [13, 14], consistent with a cardioprotective role of RAF1. BRAF is expressed in cardiomyocytes [15], but there is little information on its role in the heart.

Activating mutations in the ERK1/2 cascade can cause cancer. With oncogenic mutations in BRAF being associated with ∼30% of all cancers [16], there has been great emphasis on developing BRAF inhibitors, some of which (e.g. dabrafenib, encorafenib) are in clinical use [17, 18]. These target the highly oncogenic BRAF^V600E/K^ mutated enzyme, but also inhibit wild-type enzymes (e.g. dabrafenib has IC_50_ values of 5.2 and 6.3 nM for BRAF and RAF1, respectively [19]). This first generation of drugs are Type 1 or 1.5 inhibitors that interact with the ATP binding site, binding to and locking the enzyme in an active conformation. These drugs revealed a “RAF paradox”, in which, rather than inhibiting the pathway, they activate ERK1/2 at low concentrations [20]. This is potentially because inhibitor binding to one partner in a RAF dimer locks the other in an active conformation and (because the inhibitor is at a non-saturating concentration) this protomer activates MKK1/2 in the presence of activated Ras. Overall, the BRAF inhibitors used for cancer could affect wild-type RAF signalling in non-cancerous cells, potentially in unpredictable ways. Indeed, our recent study showed that dabrafenib (a Type 1.5 inhibitor [19]) has an anti-fibrotic effect in hypertension-induced cardiac hypertrophy in mice [21].

Here, we show that BRAF expression is upregulated in human heart failure. Activation of cardiomyocyte BRAF was sufficient to promote hypertrophy in mice, whilst the Type 1 RAF inhibitor, SB590885, and encorafenib (in clinical use) activated ERK1/2 signalling in cardiomyocytes causing hypertrophy in cultured cells. Both drugs induced cardiac hypertrophy in mouse hearts *in vivo*. Interestingly, cardiac hypertrophy induced by activation of cardiomyocyte BRAF or these RAF inhibitors was not associated with any overt fibrosis or cell infiltration.

## Materials and Methods

### Human heart samples

Failing human tissue sample collections were obtained from patients who consented to a protocol reviewed and approved by the University of Pittsburgh Institutional Review Board. Patient information is provided in **Supplementary Table S1**. Transmural tissue at the level of the anterior papillary muscle was collected at the time of cardiac transplantation from the left ventricle of end-stage heart failure patients. Samples were collected in the operating room and transported in ice-cold St. Thomas’ cardioplegia solution, flash frozen within 20 minutes of excision, and stored at -80°C prior to utilization. Non-failing heart samples were collected under University of Pittsburgh CORID #451 (Committee for Oversight of Research and Clinical Training Involving Decedents) and with consent being obtained by the local Organ Procurement Organization (OPO), CORE (Center for Organ Recovery and Education). Control left ventricular tissues were collected from hearts that were rejected for transplant for varying reasons. Tissues were collected and stored in a similar manner as the failing hearts with between 20-45 minutes of time elapsing between cross-clamp and freezing of the tissue.

### Bioinformatics analysis for transcript expression in dilated cardiomyopathy

mRNA expression of *BRAF* (ENSG00000157764), *ARAF* (NSG00000078061) and *RAF1* (ENSG00000132155) in control and diseased human hearts was determined using a previously published RNASeq dataset derived from left ventricular samples of 97 patients with end-stage dilated cardiomyopathy taken at the time of transplantation or left ventricular assist device implantation, together with 108 non-diseased controls, as described in Heinig *et al*. 2017 [22]. Differential expression analysis was carried out using DESeq2 (V*1*.*18*.*1*, Wald test) [23] between dilated cardiomyopathy patient samples and control samples.

### Ethics statement for animal experiments

Animals were housed at the BioResource Unit at University of Reading (UK registered with a Home Office certificate of designation). All procedures were performed in accordance with UK regulations and the European Community Directive 86/609/EEC for animal experiments. Work was undertaken in accordance with local institutional animal care committee procedures (University of Reading) and the U.K. Animals (Scientific Procedures) Act 1986.

### Animal housing, husbandry and welfare

Adult mice and rats were allowed to acclimatise for at least 7 d prior to experiments. Neonatal rats were culled by cervical dislocation for which additional approval and licences are not required according to UK regulations. Mice were housed in Tecniplast IVC cages (total area 512 cm^2^; maximum 5 mice per cage). Adult male rats were housed in open top NKP cages (total area 1632cm^2^; maximum 5 rats per cage). Cages were supplied with aspen sawdust bedding, sizzle nesting, cardboard tunnels and housing. Additional enrichment included chew sticks and millet to encourage foraging behaviour. Animals were provided with water and food (SDS Rm3 pelleted food for mice; SDS RM3 expanded pelleted food for rats) *ad libitum*, with a 12:12 light/dark cycle and room temperature of 21°C. Breeding was conducted with mice between 6 weeks and 8 months with a maximum of 6 litters per female. All animals were checked at least once a day by a trained, competent person and licence holders informed of any welfare issues, with consultation with a Named Veterinary Surgeon when necessary. Mice undergoing procedures were monitored using a score sheet and routinely culled if they reached a predefined endpoint agreed with the Named Veterinary Surgeon. Weights were taken before, during and at the end of the procedures. Mouse weights from the start and end of procedures are provided in **Supplementary Table S2**.

Allocation of mice to specific groups was on a random basis with randomisation performed independently of the individual leading the experiment. Mice were excluded after randomisation only if there was a health problem unrelated to the experiment, in which case they were culled. One mouse (Cre^MCM/-^, vehicle treated) was culled because of wounds inflicted as a result of aggressive behaviour. Individuals conducting the *in vivo* studies were not blinded to experimental conditions for welfare monitoring purposes. Data and sample analysis (e.g. echocardiography, histology) was performed by individuals who were blinded to intervention.

### *In vivo* mouse studies for cardiomyocyte-specific activation of BRAF

Genetically-modified mice were from Jackson Laboratories, imported into the UK and transported to University of Reading for breeding in-house. We used mice with a floxed cassette for Cre-induced knock-in of the BRAF^V600E^ mutation (B6.129P2(Cg)-*Braf* ^*tm1Mmcm*^/J, strain no. 017837) [24]. Mice were backcrossed against the C57Bl/6J background at University of Reading for at least 4 generations prior to experimentation. Mice were bred with *Myh6*-MERCreMER mice expressing tamoxifen-inducible Cre recombinase under control of the mouse *Myh6* promoter [Tg(Myh6-cre)1Jmk/J, strain no. 009074] [25]. Breeding protocols were used to produce BRAF^V600E/WT^/Cre^MCM/-^ mice, heterozygous for the BRAF^V600E^ allele and hemizygous for Cre. Cre^MCM/-^ mice, hemizygous for Cre alone, were derived from crosses between the Tg(Myh6-cre)1Jmk/J and B6.129P2(Cg)-*Braf* ^*tm1Mmcm*^/J, and treated with/without tamoxifen to control for tamoxifen-induced Cre activation.

DNA for genotyping was extracted from ear clips using Purelink genomic DNA (gDNA) mini-kits (Invitrogen) according to the manufacturer’s instructions. Briefly, tissue was digested in genomic digestion buffer containing proteinase K (overnight, 55°C). Following centrifugation (12,000 × g, 3 min, 18°C), supernatants were incubated with RNAse A (2 min) before addition of genomic lysis binding buffer mixed with an equal volume of ethanol. gDNA was purified using Purelink spin columns and PCR amplified with specific primers (**Supplementary Table S3**) using GoTaq Hot Start Polymerase (Promega). PCR conditions were 95°C for 3 min, followed by up to 33 cycles of 95°C denaturation for 30 s, 30 s annealing, elongation at 72°C for 30 s, followed by a 7 min 72°C final extension. PCR products were separated on 2% (w/v) agarose gels (25 min, 80 V) and visualised under UV light.

Male mice (8-9 wks) were treated with a single dose of tamoxifen (40 mg/kg i.p.; Sigma-Aldrich) or vehicle. Tamoxifen was dissolved in 0.25 ml ethanol and then mixed with 4.75 ml corn oil. For confirmation of recombination in the heart, RNA was extracted from heart powders and cDNA prepared as described below. cDNA (4 µl) was subjected to PCR analysis using GoTaq Hot Start Polymerase with specific primers and conditions (**Supplementary Table S3**). PCR conditions were 95°C for 5 min, followed by 40 cycles of 95°C denaturation for 40 s, 40 s annealing, elongation at 72°C for 60 s, followed by a 7 min 72°C final extension. 50% of the resulting product was digested with XbaI (New England Biolabs) (3 h, 37°C). Digested and undigested products were separated by electrophoresis on a 2% (w/v) agarose gel (85 V, 45 min). BRAF^V600E^ knock-in introduces a novel Xba1 site in the PCR product in the cDNA producing additional products [24].

### *In vivo* mouse studies for effects of SB590885 and encorafenib

Male wild-type C57Bl/6J mice (7-8 weeks of age) were from Charles River (UK). Drug delivery used Alzet osmotic pumps (models 1007D or 1004; supplied by Charles River), filled according to the manufacturer’s instructions in a laminar flow hood using sterile technique. Mice were treated with vehicle only [DMSO/PEG mix: 50% (v/v) DMSO, 20% (v/v) polyethylene glycol 400, 5% (v/v) propylene glycol, 0.5% (v/v) Tween 80 made up to 100% with H_2_O], or 0.5 mg/kg/d SB590885 (Selleck Chemicals) or 3 mg/kg/d encorafenib (MedChemExpress) dissolved in DMSO/PEG mix. Minipumps were incubated overnight in sterile PBS (37°C) prior to implantation. Implantation was performed under continuous inhalation anaesthesia using isoflurane (induction at 5%, maintenance at 2 - 2.5%) mixed with 2 l/min O_2_. A 1 cm incision was made in the mid-scapular region and mice were given 0.05 mg/kg (s.c.) buprenorphine (Ceva Animal Health Ltd.) to repress post-surgical discomfort. Minipumps were implanted portal first in a pocket created in the left flank region of the mouse. Wound closure used a simple interrupted suture with polypropylene 4-0 thread (Prolene, Ethicon). Mice were allowed to recover singly and returned to their home cage once fully recovered.

### Cardiac ultrasound

Echocardiography was performed on anaesthetised mice using a Vevo 2100 imaging system equipped with a MS400 18-38 MHz transducer (Visualsonics). Mice were anaesthetised in an induction chamber with isoflurane (5% flow rate) with 1 l/min O_2_ then transferred to the heated Vevo Imaging Station. Anaesthesia was maintained with 1.5% isoflurane delivered via a nose cone. Baseline scans were taken prior to experimentation (−7 to -3 days). Further scans were taken at intervals following tamoxifen treatment or minipump implantation. Imaging was completed within 20 min. Mice were recovered singly and transferred to the home cage once fully recovered. Left ventricular cardiac dimensions were assessed from short axis M-mode images with the axis placed at the mid-level of the left ventricle at the level of the papillary muscles. Data analysis was performed using VevoLAB software (Visualsonics) and data were gathered from two M-mode scans at each time point, taking mean values across 4 cardiac cycles. Cardiac function and global longitudinal strain were measured from B-mode long axis images using VevoStrain software for speckle tracking. Mice were culled by CO_2_ inhalation followed by cervical dislocation. Hearts were excised quickly, washed in PBS, dried and snap-frozen in liquid N_2_ or fixed for histology.

### Histology and assessment of myocyte size and fibrosis

Histological staining and analysis were performed as previously described [26], assessing general morphology by haematoxylin and eosin (H&E) and fibrosis by Masson’s trichrome and picrosirius red (PSR).

### Adult rat heart perfusions

Adult male (300-350 g) Sprague–Dawley rats were anaesthetised with a lethal intraperitoneal dose of pentobarbital sodium (60 mg/kg). After complete anaesthesia was induced, heparin (1000 U/kg) was administered intravenously. The chest cavity was opened and the heart and lungs were removed into modified ice-cold KHBBS (25 mM NaHCO_3_, 119 mM NaCl, 35 mM KCl, 2.5 mM CaCl_2_, 1.2 mM MgSO_4_, 1.2 mM KH_2_PO_4_ equilibrated with 95% O_2_/5% CO_2_) whilst the heart was still beating. Hearts were perfused retrogradely (37°C at 70 mm Hg) as described previously [27] with a 15 min equilibration period. SB590885 was added at the end of the equilibration period and all perfusions continued for a further 15 min. Hearts were ‘freeze-clamped’ between aluminium tongs cooled in liquid nitrogen and pulverized under liquid N_2_ in a pestle and mortar. The powders were stored at −80°C.

### Cardiomyocyte cultures

Neonatal rat ventricular myocytes were prepared and cultured from 2-4 d Sprague-Dawley rats (Charles River) as previously described [28]. Briefly, neonatal rats were culled by cervical dislocation, then the ventricles were dissected and dissociated by serial digestion at 37°C with 0.44 mg/ml (6800 U) Worthington Type II collagenase (supplied by Lonza) and 0.6 mg/ml pancreatin (Sigma-Aldrich, cat. No. P3292). Cell suspensions were recovered by centrifugation (5 min, 60 × g) and resuspended in plating medium (Dulbecco’s modified Eagle’s medium (DMEM)/medium 199 [4:1 (v/v)]) containing 15% (v/v) foetal calf serum (FCS; Life Technologies) and 100 U/ml penicillin and streptomycin. Cells were pre-plated on plastic tissue culture dishes (30 min) to remove non-cardiomyocytes. Non-adherent cardiomyocytes were collected and viable cells counted by Trypan Blue (Sigma-Aldrich) exclusion using a haemocytometer. For immunoblotting or qPCR, viable cardiomyocytes were plated at a density of 4 × 10^6^ cells/dish on 60 mm Primaria dishes pre-coated with sterile 1% (w/v) gelatin (Sigma-Aldrich). After 18 h, myocytes were confluent and beating spontaneously. For immunostaining experiments, cardiomyocytes were plated at 1 × 10^6^ cells/dish on 35 mm Primaria dishes containing glass coverslips pre-coated with sterile 1% (w/v) gelatin followed by laminin (20 µg/ml in PBS; Sigma-Aldrich). After 18 h, myocytes were ∼50% confluent and beating spontaneously. For all experiments, the plating medium was withdrawn after 18 h and cells were incubated in serum-free maintenance medium (DMEM/medium 199 [4:1 (v/v)] containing 100 U/ml penicillin and streptomycin) for a further 24 h prior to experimentation.

### RNA preparation and qPCR

Total RNA was prepared using RNA Bee (AMS Biotechnology Ltd) with 1 ml per 4 × 10^6^ cardiomyocytes or 10-15 mg mouse heart powder. RNA was prepared according to the manufacturer’s instructions, dissolved in nuclease-free water and purity assessed from the A_260_/A_280_ measured using an Implen NanoPhotometer (values of 1.8–2.1 were considered acceptable). RNA concentrations were determined from the A_260_ values. Quantitative PCR (qPCR) was performed as previously described [28]. Total RNA (1 µg) was reverse transcribed to cDNA using High Capacity cDNA Reverse Transcription Kits with random primers (Applied Biosystems) according to the manufacturer’s instructions. qPCR was performed using an ABI Real-Time PCR 7500 system (Applied Biosystems). Optical 96-well reaction plates were used with iTaq Universal SYBR Green Supermix (Bio-Rad Laboratories Inc.) according to the manufacturer’s instructions. Primers were from PrimerDesign, Eurofins Genomics or Thermo Fisher Scientific (see **Supplementary Table S4**). *GAPDH* was used as the reference gene for the study. Results were normalized to *GAPDH*, and relative quantification was obtained using the ΔCt (threshold cycle) method; relative expression was calculated as 2^−ΔΔCt^, and normalised to vehicle or time 0.

### Immunoblotting

Human heart samples were homogenised in RIPA buffer (Thermo Scientific, Cat. No. 89901) supplemented with Complete Mini, EDTA-free protease inhibitors (Roche, Cat. No. 11836170001) and PhosSTOP easypack phosphatase inhibitors (Roche, Cat. No. 04906837001). Samples were centrifuged (3,000 × g, 15 min, 4°C) and the supernatants collected. Protein quantification was performed using a Pierce BCA Protein Assay kit (Thermo Scientific, Cat. No. 23225). Samples were boiled with 0.33 vol SDS-polyacrylamide gel electrophoresis (SDS-PAGE) sample buffer [0.33 M Tris-HCl pH 6.8, 10% (w/v) SDS, 13% (v/v) glycerol, 133 mM dithiothreitol, 0.2 mg/ml bromophenol blue].

Rodent hearts were ground to powder under liquid N_2_. Samples (15-20 mg) were extracted in 8 vol (relative to powder weight) Buffer A [20 mM β-glycerophosphate (pH 7.5), 50 mM NaF, 2 mM ethylenediamine tetraacetic acid (EDTA), 1% (v/v) Triton X-100, 5 mM dithiothreitol] containing protease and phosphatase inhibitors [10 mM benzamidine, 0.2 mM leupeptin, 0.01 mM trans-epoxy succinyl-l-leucylamido-(4-guanidino)butane, 0.3 mM phenylmethylsulphonyl fluoride, 4 µM microcystin]. For total cell extracts, cells were washed with ice-cold PBS and scraped into 150 µl Buffer A containing protease and phosphatase inhibitors. Samples were vortexed and extracted on ice (10 min). Extracts were centrifuged (10,000 × g, 10 min, 4°C). The supernatants were removed, a sample was taken for protein assay and the remainder boiled with 0.33 vol sample buffer. Protein concentrations were determined by BioRad Bradford assay using bovine serum albumin (BSA) standards.

For analysis of cytosolic and nuclear extracts, cells were washed in ice-cold PBS and harvested into 150 µl of hypotonic buffer (10 mM Hepes, pH 7.9, 10 mM KCl, 1.5 mM MgCl_2_, 0.3 mM Na_3_VO_4_) containing protease and phosphatase inhibitors. Samples were centrifuged (10,000 × g, 4°C, 5 min) and the supernatants (cytosolic fractions) boiled with 50 µl of sample buffer. Pellets were resuspended in 50 µl of nuclear extraction buffer (20 mM Hepes pH 7.9, 420 mM NaCl, 1.5 mM MgCl_2_, 0.2 mM EDTA, 25% (v/v) glycerol, 0.3 mM Na_3_VO_4_) containing protease and phosphatase inhibitors, and extracted on ice for 1 h with occasional vortex-mixing. The samples were centrifuged (10,000 × g, 4°C, 5 min) and the supernatant nuclear extracts boiled with 20 µl sample buffer.

Proteins were separated by SDS-PAGE on 10% (w/v) polyacrylamide resolving gels with 6% stacking gels, transferred electrophoretically to nitrocellulose using a BioRad semi-dry transfer cell (10 V, 60 min) and detected as previously described [28]. Bands were visualised by enhanced chemiluminescence using ECL Prime Western Blotting detection reagents and using an ImageQuant LAS4000 system (GE Healthcare). ImageQuant TL 8.1 software (GE Healthcare) was used for densitometric analysis. Raw values for phosphorylated kinases were normalised to the total kinase. Values for all samples were normalised to the mean of the controls. Details of all primary and secondary antibodies are in **Supplementary Table S5**.

### Immunostaining

Cardiomyocytes were washed with ice-cold PBS and fixed in 3.7% (v/v) formaldehyde in PBS (10 min, room temperature). Cells were permeabilised with 0.3% (v/v) Triton X-100 (10 min, room temperature) in PBS, and non-specific binding blocked with 1% (w/v) fatty acid-free BSA (Sigma-Aldrich UK) in PBS containing 0.1% (v/v) Triton X-100 (10 min, room temperature). All incubations were at room temperature in a humidified chamber, and coverslips were washed three times in PBS after each stage of the immunostaining procedure. Cardiomyocytes were stained with mouse monoclonal primary antibodies to troponin T (60 min) with detection using anti-mouse immunoglobulin secondary antibodies coupled to Alexa-Fluor 488 (60 min). Details of antibodies are in **Supplementary Table S5**. Coverslips were mounted using fluorescence mounting medium (Dako) and viewed with a Zeiss Axioskop fluorescence microscope using a 40× objective. Digital images were captured using a Canon PowerShot G3 camera using a 1.8× digital zoom and cardiomyocyte sizes measured using ImageJ. Images were cropped for presentation using Adobe Photoshop CC 2018.

### Fast protein liquid chromatography (FPLC), Raf kinase immunoprecipitation and protein kinase assays

Protein complexes were separated according to relative molecular mass using an Äkta™ FPLC system with a Superdex 200 HR 10/30 column (GE Healthcare) and 24 × 10^6^ cardiomyocytes per sample. Fractions (equal volumes) were immunoblotted as described above. The column was calibrated using gel filtration calibration kits (GE Healthcare). BRAF or RAF1 was immunoprecipitated and assayed with GST-MKK1 as substrate, largely as described in [29] using 1 µg of antibody per sample and 4 × 10^6^ NRVMs per sample. Samples of total extracts, immunoprecipitates and supernatants were boiled in sample buffer for immunoblotting with antibodies selective for phosphorylated or total MKK1/2, BRAF or RAF1.

### Statistical analysis

Data analysis used Microsoft Excel and GraphPad Prism 8.0. Statistical analysis was performed using GraphPad Prism 8.0 with two-tailed unpaired t tests, or two-tailed one-way or two-way ANOVA as indicated in the Figure Legends. A multiple comparison test was used in combination with ANOVA. A Grubb’s outlier test was applied to the data, and outliers excluded from the analysis. Graphs were plotted with GraphPad Prism 8.0. Specific p values are provided with significance levels of p<0.05 in bold type.

## Results

### BRAF is upregulated in human failing hearts

BRAF and RAF1 (but not ARAF) mRNA (**Figure 1A**) and protein (**Figure 1B,C**)were significantly upregulated in samples from a cohort of 12 patients with heart failure of mixed non-ischaemic aetiology compared with normal controls. Since this group is relatively small, we mined an RNASeq database of patients with well-defined dilated cardiomyopathy (n=97) *vs* normal controls (n=108) [22]. *ARAF, BRAF* and *RAF1* transcripts were readily detected in human hearts, but only *BRAF* transcript expression was significantly increased in dilated cardiomyopathy samples; *ARAF* and *RAF1* transcripts showed decreased expression (**Figure 1D**). Thus, BRAF is potentially important in human heart failure.

**Figure 1.**
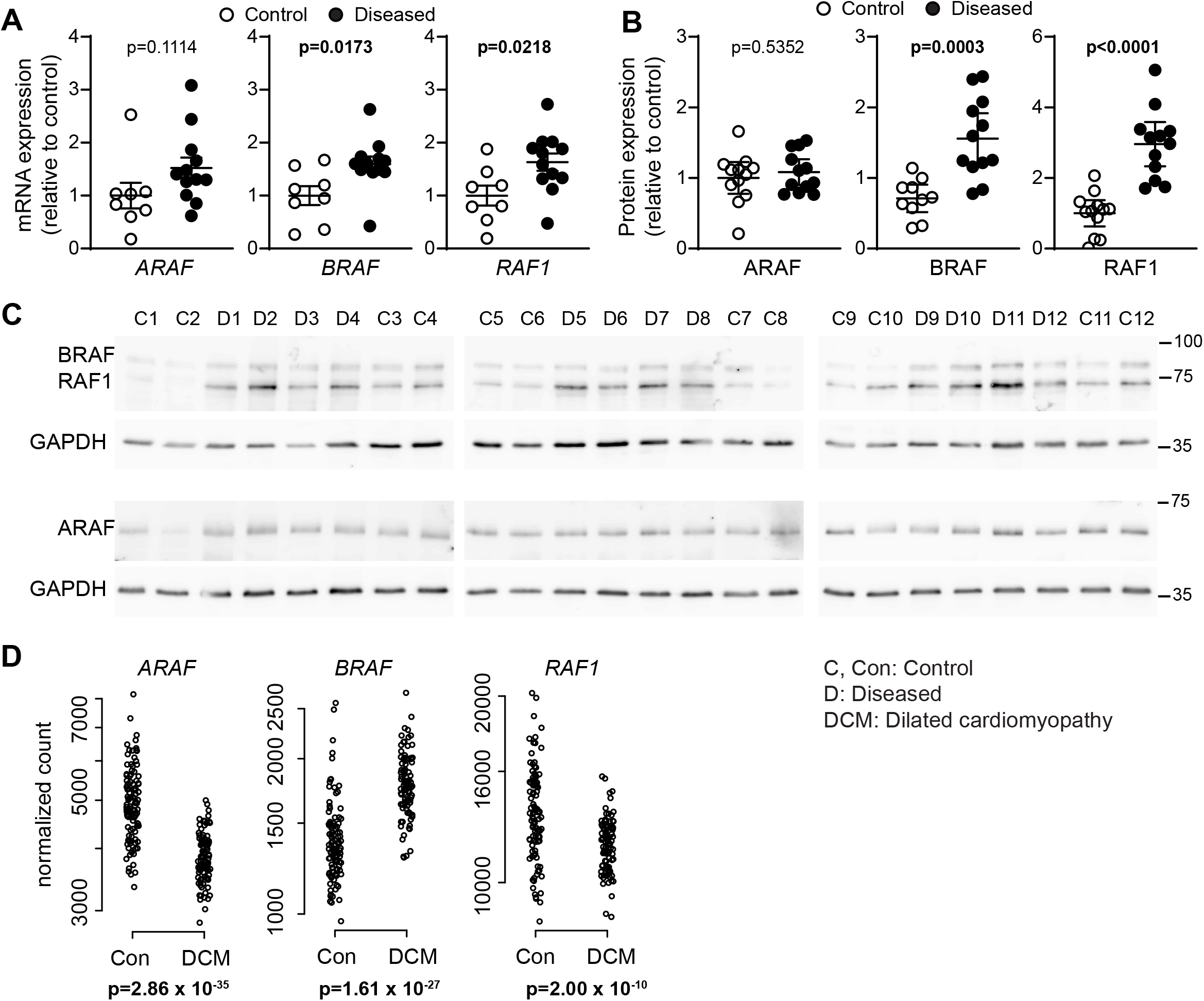
BRAF is upregulated in human failing hearts. **A-C**, Expression of ARAF, BRAF and RAF1 in human heart samples showed increased expression of BRAF and RAF1 in failing hearts compared with controls (unpaired two-tailed t tests). **A**, mRNA expression was assessed by qPCR. **B-C**, Proteins were analysed by immunoblotting. Blots were reprobed for GAPDH. Densitometric analysis of the blots (**C**) is in (**B**). Positions of relative molecular mass markers (kDa) are on the right of the immunoblots. Data were normalised to GAPDH and expressed relative to the mean of the controls. **D**, Data were mined from an RNASeq database of patients with dilated cardiomyopathy (n=97) and normal controls (n=108). Data for individual samples are shown with adjusted p values.

### Activation of cardiomyocyte BRAF promotes hypertrophy

To assess the role of BRAF in cardiomyocytes, we used a genetic model with tamoxifen-inducible Cre regulated by a *Myh6* promoter for cardiomyocyte-specific expression (Cre^MCM^) [25]. This allowed selective manipulation of BRAF activity in adult cardiomyocytes, avoiding complications during development. BRAF was activated in cardiomyocytes using a floxed gene for BRAF^V600E^ knock-in into the endogenous gene [24]. We used mice heterozygous for the floxed gene for BRAF^V600E^ knock-in and hemizygous for Cre (i.e. BRAF^V600E/WT^/Cre^MCM/-^). Following baseline echocardiography, recombination was induced with a single injection of 40 mg/kg tamoxifen, a dose established to induce recombination in the absence of significant cardiomyopathy [30], and an approach which allows rapid clearance from the body (within 24-72 h) [31]. Echocardiograms were taken and mice sacrificed at intervals as shown in the scheme in **Figure 2A**. Tamoxifen-treatment induced selective recombination in the heart (**Figure 2B**). Recombination in heart tissue is not complete because approximately 70% of cells in the heart are cardiac non-myocytes (e.g. endothelial cells and fibroblasts). At 24 h, there was no change in expression of BRAF mRNA expression, although there was a small, significant increase and decrease in *ARAF* and *RAF1* mRNA expression, respectively (**Figure 2C**). This had normalised by 10 d, with no significant effects on expression of any of the RAF kinases at the mRNA or protein level (**Figure 2C,D**). BRAF^V600E^ knock-in increased phosphorylation (i.e. activation) of MKK1/2 over 10 d (**Figure 2D**), with upregulation of mRNAs for ERK1/2-dependent immediate early genes associated with cardiac hypertrophy (e.g. *Egr1, Dusp5, Fos, Jun* and *Myc*) [28, 32, 33], along with hypertrophic gene markers (*Nppa, Nppb, Myh7*) [2], consistent with cardiac hypertrophy (**Figure 2E,F**). There was also a small, but significant increase in *IL11* mRNA expression (known to drive ERK1/2-dependent fibrosis in the heart [34]) that, together with *Col1a1* and *Lox*, suggested there could be some fibrotic response. A small, but significant increase in *IL1β* was detected suggesting there could be some inflammation. Tamoxifen alone induced a small increase in mRNA expression of *Myh7* in hearts of mice hemizygous for Cre only, but did not affect the other genes (**Supplementary Table S6**).

**Figure 2.**
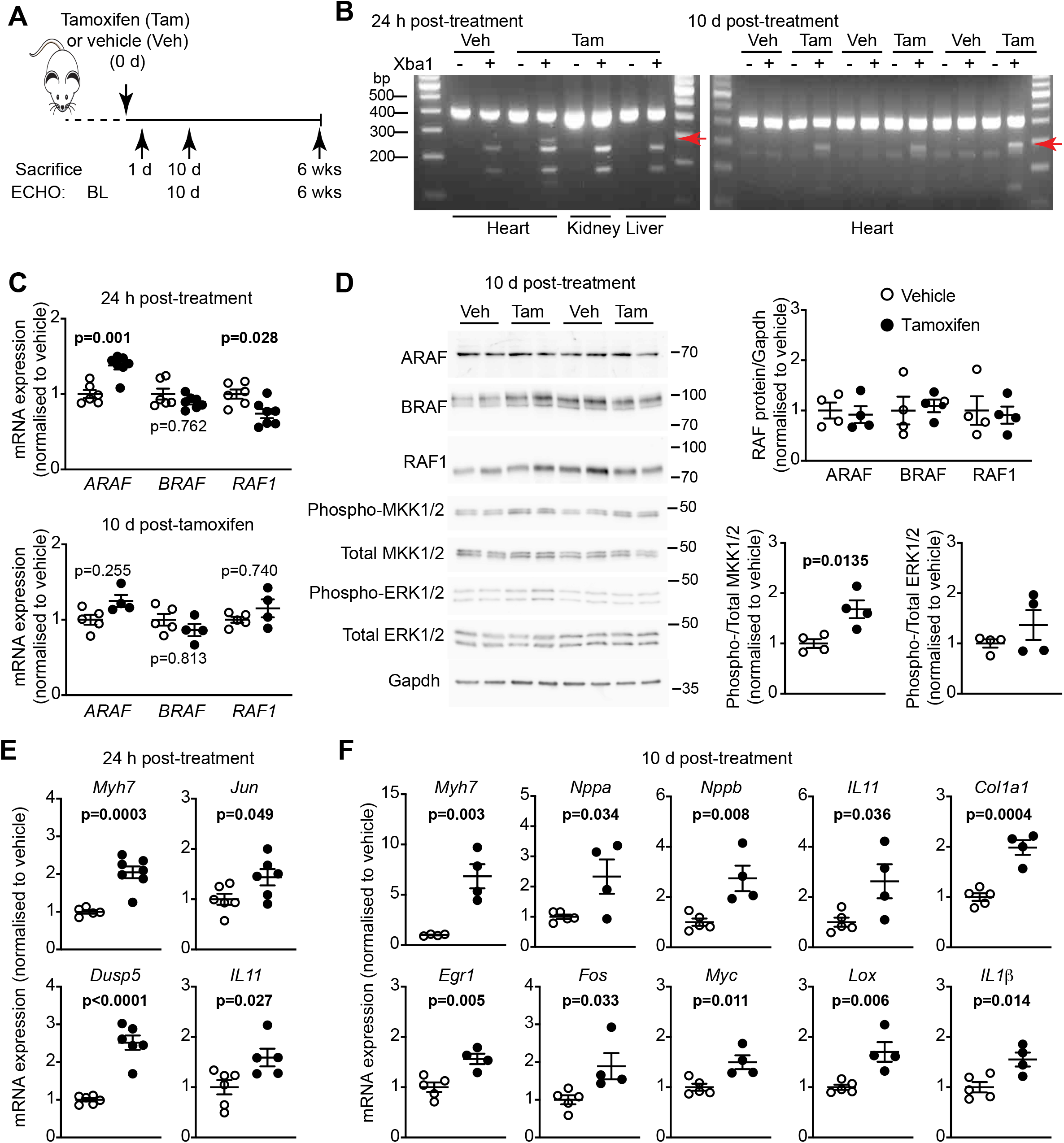
Activated cardiomyocyte BRAF induces changes in cardiac gene expression in mouse hearts *in vivo*. **A**, Study protocol. Mice heterozygous for floxed BRAF^V600E^ and hemizygous for Myh6-directed tamoxifen-inducible Cre were treated with 40 mg/kg (i.p.) tamoxifen (Tam) or vehicle (Veh). Echocardiograms (ECHO) were taken at baseline (BL) then at 10 d or 6 weeks after tamoxifen treatment. **B**, Confirmation of recombination for BRAF^V600E^ knock-in was based on introduction of an Xba1 restriction site in the mRNA following recombination. cDNA was prepared from RNA extracted from heart, liver or kidney, 24 h or 10 d after treatment with tamoxifen or vehicle. cDNA was PCR amplified and subject to Xba1 digestion, resulting in appearance of additional bands (red arrows) in hearts (not liver or kidney) following treatment with tamoxifen, but not vehicle. **C**, qPCR analysis of mRNA expression of RAF isoforms in samples of mouse hearts treated with vehicle or tamoxifen for 24 h (upper panel) or 10 d (lower panel). **D**, Immunoblotting of RAF isoforms, phosphorylated (phospho-) or total MKK1/2, and phosphorylated (phospho-) or total ERK1/2 in samples of mouse hearts treated with vehicle or tamoxifen (10 d). Representative immunoblots are on the left (positions of relative molecular mass markers in kDa are on the right of each blot) with densitometric analysis on the right. **E**,**F**, qPCR for mRNA expression in mouse hearts after tamoxifen-treatment for 24 h (**E**) or 10 d (**F**). Data are provided as individual values with means ± SEM. Statistical analysis: unpaired two-tailed t test.

We assessed the effects of activated cardiomyocyte BRAF on cardiac function and dimensions using echocardiography (**Figure 3; Supplementary Table S7**). Assessment of left ventricular dimensions from M-mode short-axis images of the heart indicated that activation of BRAF in cardiomyocytes increased diastolic and systolic wall thicknesses within 10 d of tamoxifen treatment (**Figure 3A,B**). This was associated with an increase in cardiomyocyte cross-sectional area indicative of hypertrophy (**Figure 3C**). Tamoxifen treatment of mice hemizygous for Cre did not have any significant effect on cardiac function or dimensions (**Supplementary Table S8**). The increase in left ventricular wall thickness was sustained over 6 weeks (**Figure 3D,E**). By this stage, there was a significant decrease in internal diameters, particular in systole, suggesting there was likely to be a change in cardiac function. Assessment of B-mode long-axis images of the heart at 6 weeks with strain analysis using speckle-tracking software demonstrated that activation of cardiomyocyte BRAF increased the predicted LV mass, with a reduction in diastolic and systolic volumes, increasing ejection fraction and fractional shortening (**Figure 3F; Supplementary Table S9**). Histological assessment demonstrated that the increase in cardiomyocyte cross-sectional was sustained after 6 weeks of BRAF activation (**Figure 3G**). Despite the increase in *Nppa, Nppb, IL11, Col1a1, Lox* and *IL1β* mRNAs (**Figure 2E,F**), here was no evidence for any increase in cardiac fibrosis or cellular damage at 10 d or 6 weeks (**Figure 3C,G**). We conclude that the changes in mRNA expression of these genes were either too small to elicit a significant effect overall, or that these genes were subject to translational regulation. Overall, the data indicate that activation of endogenous cardiomyocyte BRAF induces cardiac hypertrophy and enhances cardiac function.

**Figure 3.**
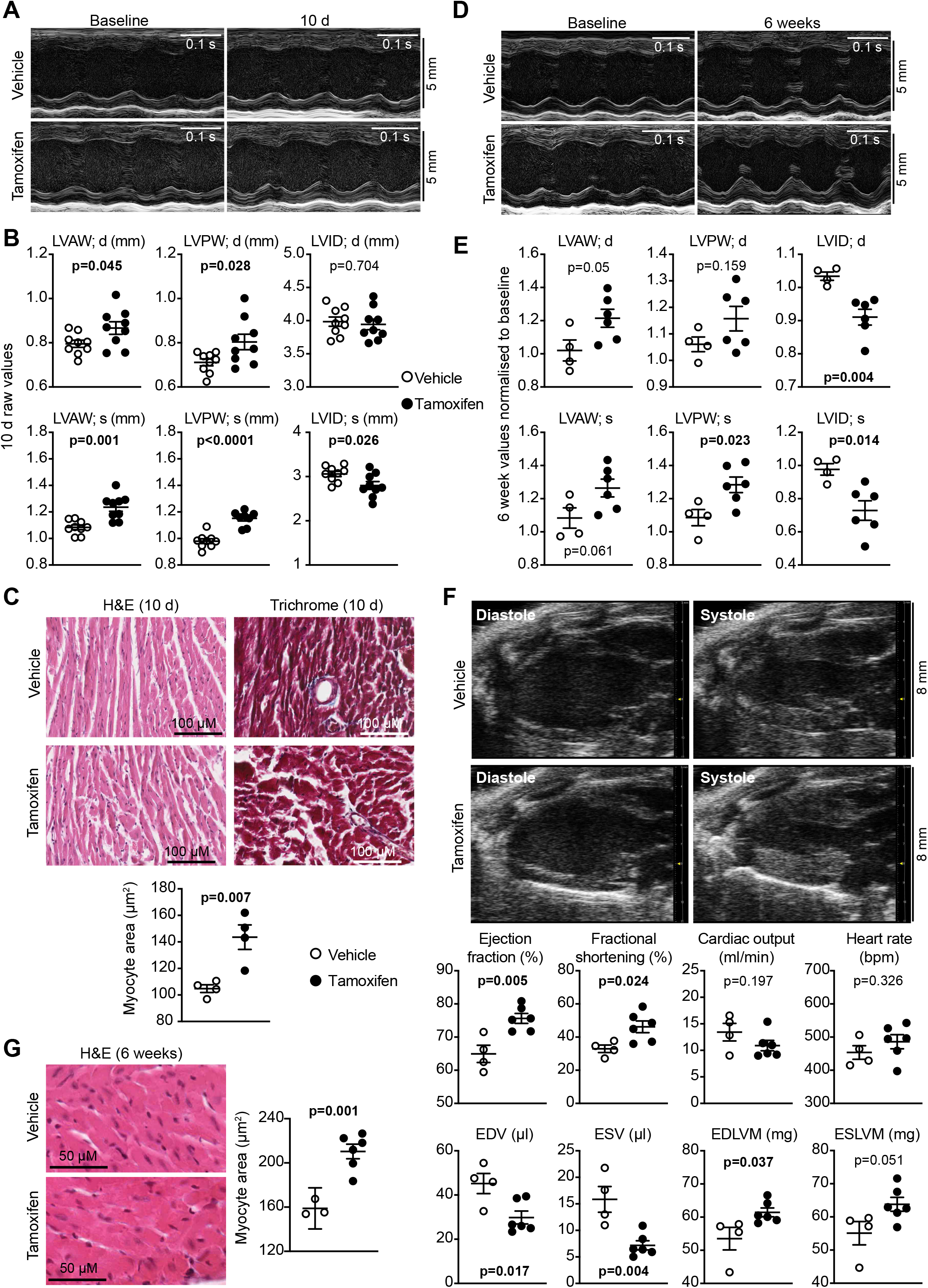
Activated cardiomyocyte BRAF promotes cardiac hypertrophy in mouse hearts *in vivo*. Mice heterozygous for floxed BRAF^V600E^ and hemizygous for Myh6-directed tamoxifen-inducible Cre (BRAF^V600E/WT^ /Cre^MCM/-^) were treated with 40 mg/kg (i.p.) tamoxifen or vehicle for 10 d (**A-C**) or 6 weeks (**D-G**). Cardiac function and dimensions were measured by echocardiography. **A** and **D**, Representative M-mode images from short-axis views for individual mice before and after treatment. **B** and **E**, Quantification of echocardiograms represented in panels **A** and **D**, respectively. LV, left ventricle; AW, anterior wall; PW, posterior wall; ID, internal diameter; d, diastole; s, systole. **C** and **G**, Sections of BRAF^V600E/WT^ /Cre^MCM/-^ mouse hearts stained with haemotoxylin and eosin (H&E) or Masson’s Trichrome were analysed. Representative images are shown with quantification provided below (**C**) or in the right panel (**G**). **F**, B-mode images of long-axis views of the heart were used for speckle-tracking strain analysis. Representative images are in the upper panels with cardiac function data in the lower panels. EDV, end diastolic volume; ESV, end systolic volume; EDLVM, end diastolic left ventricular mass; ESLVM, end systolic LV mass. Data are provided as individual points with means ± SEM. Statistical analysis: unpaired two-tailed t test.

### BRAF and RAF1 form high molecular weight complexes in cardiomyocytes and are activated by the Type 1 RAF inhibitor, SB590885

ARAF, RAF1 and BRAF have predicted relative molecular masses of 68, 73 and 85 kDa, but RAF1/BRAF operate as dimers to activate MKK1/2 [20], potentially in a multiprotein complex, and ARAF may serve as a scaffold [35]. In proliferating cells, activation of RAF signalling by “RAF paradox”-inducing compounds may be associated with dimer formation or stabilisation [20]. However, cardiomyocytes are terminally-differentiated and RAF signalling is rapid in these cells (e.g. activation of RAF1 by the pro-hypertrophic agonist endothelin-1 is maximal within 1-3 min [11]), suggesting the complexes may be pre-formed. We separated RAF complexes from cardiomyocytes using Superdex 200 gel filtration FPLC (**Figure 4A-D**). BRAF was detected largely in fractions 34-38 correlating with relative molecular mass complexes of 230-370 kDa, whereas RAF1 and ARAF eluted across a wider range corresponding to relative molecular mass complexes of 260-580 kDa. We detected little RAF in fractions 40/41 or 46-48 where dimers or monomers would be expected to elute, suggesting that RAFs predominantly exist in multi-protein complexes in cardiomyocytes. To assess BRAF and RAF1 activities, we first immunoblotted cardiomyocyte extracts with antibodies to the activating phosphorylation sites (**Figure 4E**). As shown previously [11], endothelin-1 substantially increased phosphorylation of RAF1(S338). This was associated with a band shift (appearance of bands of apparently higher molecular weight) of total RAF1 protein consistent with phosphorylation. Endothelin-1 had little effect on phosphorylation of BRAF(S445), but induced a band shift of the total protein suggesting that there was or had been activation. Using *in vitro* protein kinase assays, we determined that activation of RAF1 by endothelin-1 was substantially greater than BRAF (**Figure 4F**). RAF1 was detected in BRAF immunoprecipitates, but it was difficult to detect BRAF in RAF1 immunoprecipitates. This may indicate that RAF1 is of higher abundance and interacts with proteins other than BRAF or it could be a technical issue relating to the affinities of the different antibodies. Nevertheless, BRAF and RAF1 co-immunoprecipitated and activation by endothelin-1 did not affect the degree of association (**Figure 4F**).

**Figure 4.**
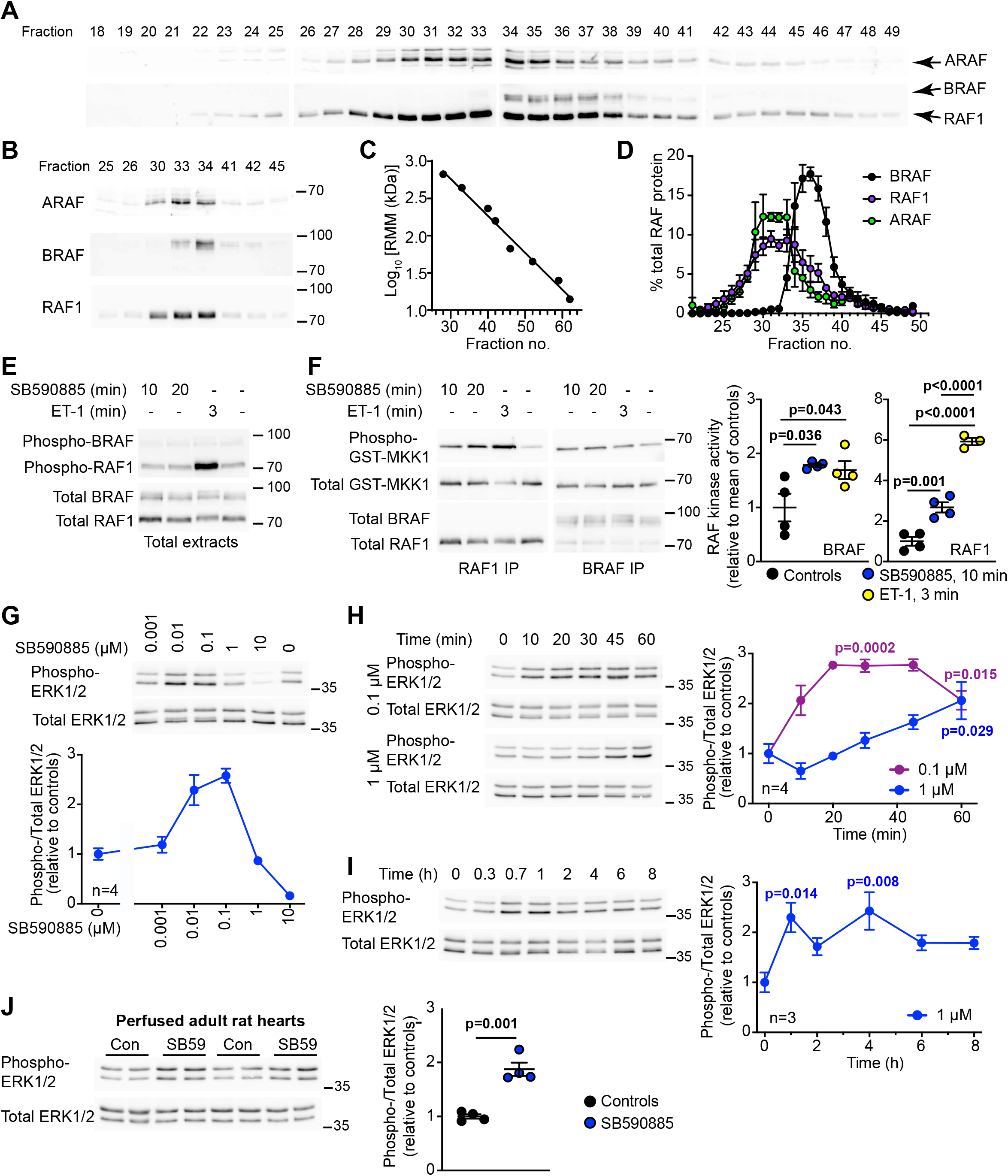
Activation of RAF and ERK1/2 signalling in cardiomyocytes by the type 1 RAF kinase inhibitor, SB590885. **A-D**. Superdex 200 FPLC separation of RAF complexes in unstimulated neonatal rat cardiomyocytes. Fractions were immunoblotted with antibodies to ARAF, or BRAF and RAF1. Representative blots of all column fractions (**A**) and comparing expression in fractions across different blots (**B**) are shown. **C**, Column calibration with relative molecular mass (RMM) standards. **D**, Elution profiles of ARAF, BRAF and RAF1. Results are means ± SEM (n=3). **E**, Immunoblots of extracts from neonatal rat cardiomyocytes exposed to 0.1 µM SB590885 or 0.1 µM endothelin-1 (ET-1) for the times indicated. Total extracts were immunoblotted for phosphorylated BRAF(S445) and RAF1(S338) (upper blot) or the total proteins (lower blot). Representative blots of 4 independent cardiomyocyte preparations are shown. **F**, RAF kinase activities in neonatal rat cardiomyocytes exposed to the indicated concentrations of SB590885 or 0.1 µM ET-1 for the times shown. Activities were assayed using GST-MKK1 following immunoprecipitation (IP) of RAF1 or BRAF. Phosphorylation of GST-MKK1 was detected with antibodies to phosphorylated MKK1/2. Assay samples were also immunoblotted for total MKK1/2, BRAF and RAF1. Representative blots are shown (left panels) with densitometric quantification (right panels; results are the ratio of phosphorylated/total GST-MKK1, normalised to the mean of controls). Data are provided for each experiment with means ± SEM (n= 4). Statistical analysis used one-way ANOVA with Holm-Sidak’s post-test. **G-J**, Immunoblot analysis of phosphorylated (phospho-) and total ERK1/2 in extracts from neonatal rat cardiomyocytes (**G, H** and **I**) or Langendorff perfused adult rat hearts (**J**). Cells were exposed to the indicated concentrations of SB590885 for 20 min (**G**) or the indicated concentration of SB590885 for the times shown (**H** and **I**). Rat hearts were perfused (15 min) under control conditions or with 1 µM SB590885 (SB59) (**J**). Representative immunoblots are shown with densitometric analysis. Results of the cell experiments are provided as means ± SEM for n=3 or 4 independent cell preparations as indicated. Heart perfusion data are provided as individual results with means ± SEM (n= 4). Statistical analysis used one-way ANOVA with Holm-Sidak’s post-test and individual p values are provided. Positions of relative molecular mass markers (kDa) are on the right of the immunoblots.

We next assessed the effects of SB590885, a Type 1 RAF kinase inhibitor with 11-fold higher affinity for BRAF than RAF1 [36], on activation of ERK1/2 in cardiomyocytes. As expected, high concentrations (10 µM) inhibited ERK1/2 phosphorylation, but concentrations of 0.1-0.01 µM SB590885 activated ERK1/2 (**Figure 4G**). Low concentrations of SB590885 also increased MKK1/2-activating activity in RAF1 or BRAF immunoprecipitates (**Figure 4F**). We assessed the time courses for ERK1/2 activation by SB590885 in cardiomyocytes, comparing 1 µM (with no effect on ERK1/2 phosphorylation at 20 min) with 0.1 µM (that activated ERK1/2). The higher concentration activated ERK1/2 at later times giving a similar degree of activation at 1 h as 0.1 µM, and ERK1/2 activation remained elevated over at least 8 h (**Figure 4H,I**). This may reflect reducing concentrations of active, available compound as it is metabolised. To confirm that the RAF paradox was relevant for the adult heart, we assessed the effects of SB590885 on ERK1/2 phosphorylation in Langendorff perfused adult rat hearts. Consistent with cardiomyocytes, SB590885 activated ERK1/2 (**Figure 4J**).

In cardiomyocytes, hypertrophic agonists (e.g. endothelin-1) increase activated ERK1/2 in the nuclear compartment without net accumulation of total ERK1/2 and this regulates gene expression [33]. SB590885 increased ERK1/2 phosphorylation in the nuclear-enriched protein fraction (**Figure 5A,B**) and increased expression of a range of immediate early genes (**Figure 5C**). The latter was inhibited by the MKK1/2 inhibitor PD184352 (2 µM) [37], confirming the response was a direct consequence of ERK1/2 signalling. Consistent with a role for ERK1/2 signalling in cardiomyocyte hypertrophy, SB590885 increased cardiomyocyte cross-sectional area (**Figure 5D**).

**Figure 5.**
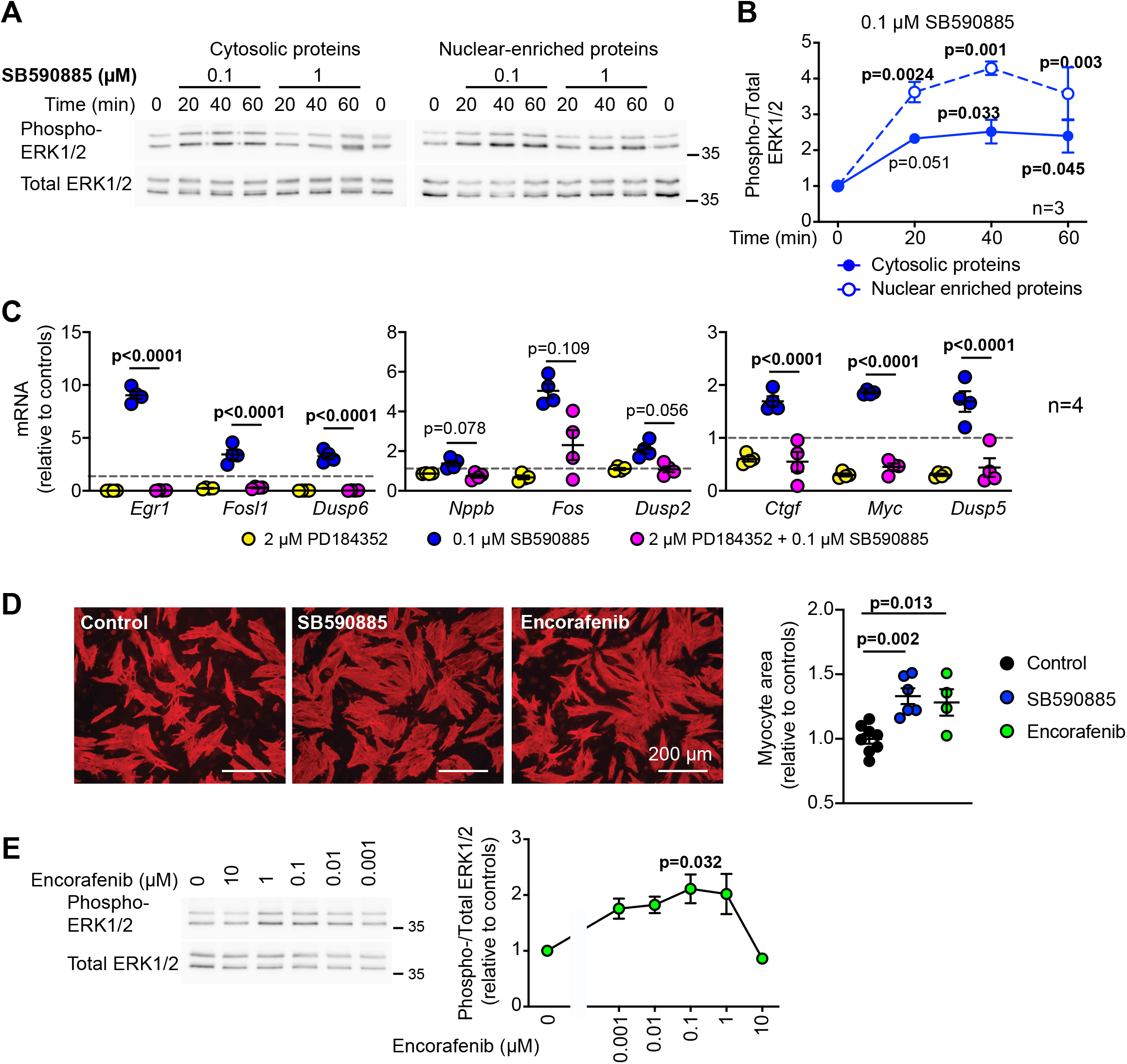
SB590885 promotes immediate early gene expression and hypertrophy in cardiomyocytes. **A** and **B**, Rat neonatal cardiomyocytes were exposed to the indicated concentrations of SB590885 for the times shown. Cytosolic and nuclear-enriched fractions were immunoblotted for phosphorylated (phospho-) or total ERK1/2. Representative blots are shown (**A**) with quantification for 0.1 µM SB590885 (**B**). Results are means ± SEM (n=3). p values are relative to 0 min (one-way ANOVA with Holm-Sidak’s post-test). Positions of relative molecular mass markers (kDa) are on the right of the immunoblots. **C**, Rat neonatal cardiomyocytes were untreated or exposed to 0.1 µM SB590885 (1 h) with/without 2 µM PD184352. mRNA expression was measured by quantitative qPCR. Individual data points are shown with means ± SEM (n=4). p values are for PD184352/SB590885 *vs* SB590885 (one-way ANOVA with Holm-Sidak’s post-test). **D**, Cardiomyocytes were incubated with 1 µM SB590885, 1 µM encorafenib or under control conditions (24 h) and immunostained for troponin T. Images (left panels) are representative of 6 (SB590885, controls) or 4 (encorafenib) independent myocyte preparations with analysis of cell size shown in the right panel. Images are from the same experiment. Individual data points are shown with means ± SEM. Statistical analysis used a two-tailed paired t test. **E**, Immunoblot analysis of phosphorylated (phospho-) and total ERK1/2 in extracts from neonatal rat cardiomyocytes exposed to the indicated concentrations of encorafenib (20 min). Representative blots are shown on the left with densitometric analysis on the right. Results are means ± SEM (n=3 independent cell preparations). p value is relative to controls with no encorafenib (one-way ANOVA with Holm-Sidak’s post-test).

SB590885 was developed as a potential therapeutic agent but is not used clinically. Our data suggest it may not be particularly stable even in cells in culture (**Figure 4H,I**) so we considered if other RAF inhibitors may exert a similar effect. Of the inhibitors in clinical use, encorafenib is available as an anti-cancer therapy for use in combination with the MKK1/2 inhibitor binimetinib [38]. It is not clear if encorafenib is a Type 1 RAF inhibitor, but it activated ERK1/2 signalling in cultured cardiomyocytes at low concentrations (0.001 – 1 µM) (**Figure 5E**), and promoted an increase in cardiomyocyte cross-sectional area (**Figure 5D**).

### SB590885 and encorafenib promote cardiomyocyte and cardiac hypertrophy in vivo

Our studies of SB590885 and encorafenib in cardiomyocytes suggested they should be pro-hypertrophic *in vivo*. To determine if this is the case, we treated adult male mice with SB590885 using osmotic minipumps for drug delivery. We used echocardiography to assess the effects on cardiac dimensions (from M-mode images of short-axis views of the heart) and function (by strain analysis of B-mode images of the long-axis views of the heart). SB590885 is not used clinically, so we selected a concentration based on the dose of the alternative RAF inhibitor, dabrafenib. Dabrafenib is administered orally in humans at up to 150 mg, twice daily (i.e. ∼3-5 mg/kg/d). IC_50_ values for inhibition of BRAF and RAF1 by dabrafenib are 5.2 nM and 6.3 nM, respectively [19]. IC_50_ values for SB590885 for inhibition of BRAF and RAF1 are ∼6-fold less than dabrafenib for inhibition of RAF1 [36], so we used 0.5 mg/kg/d. SB590885 promoted a small increase in diastolic and systolic left ventricular wall thickness within 3 d (**Figure 6A; Supplementary Table S10**). This was associated with a significant increase in predicted left ventricular mass (**Figure 6B; Supplementary Table S9**). Consistent with the echocardiography, SB590885 induced an increase in cardiomyocyte cross-sectional area without any overt increase in fibrosis or infiltrating cells (**Figure 6C**). However, in contrast to BRAF^V600E^ knock-in (**Figure 2E,F**), there were no significant changes in mRNA expression of *Nppa* or *Nppb* (**Figure 6D**).

**Figure 6.**
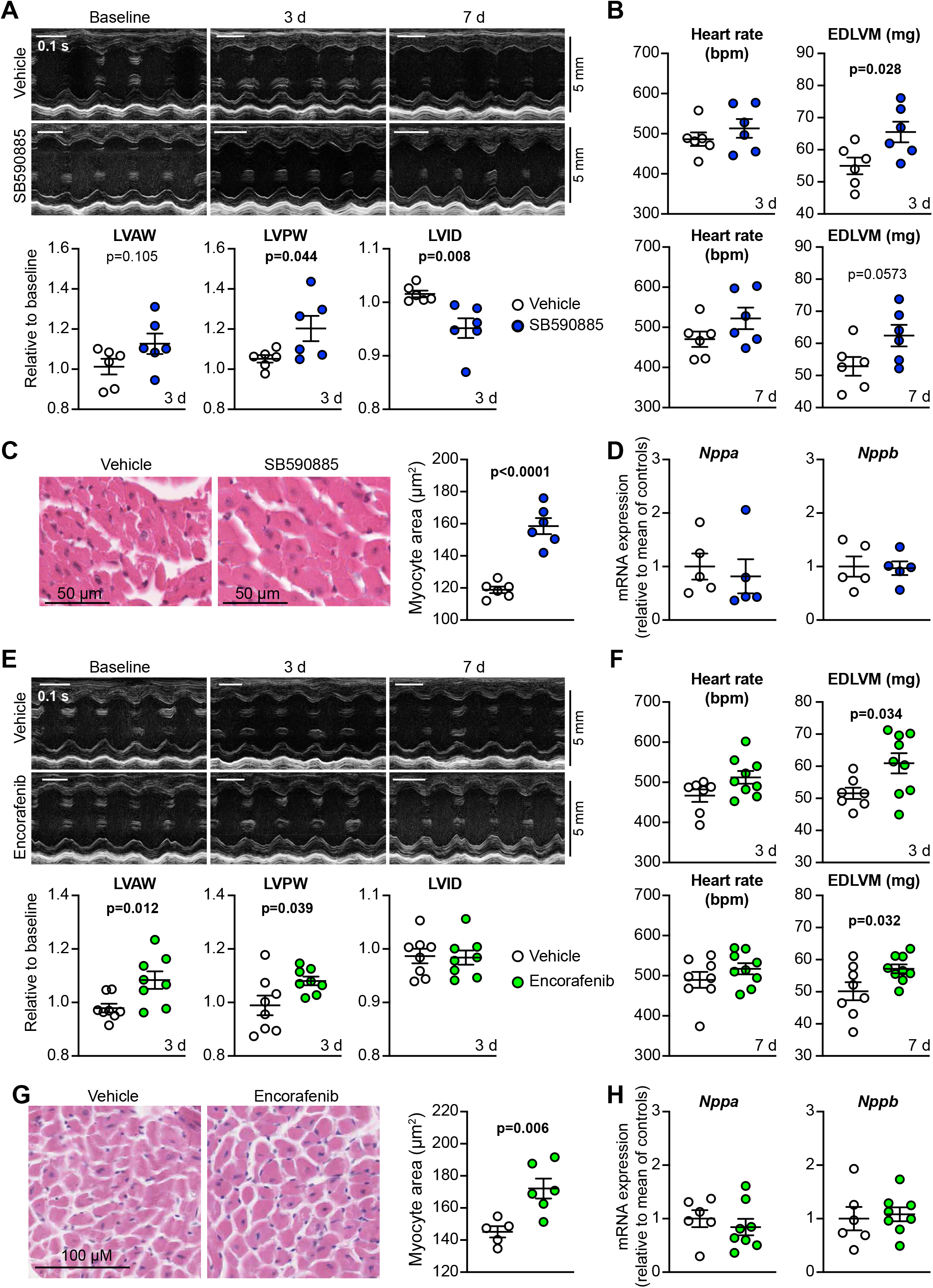
SB590885 and encorafenib promote cardiac hypertrophy *in vivo*. C57Bl/6J mice were treated with vehicle, 0.5 mg/kg/d SB590885 (**A-D**), or 3 mg/kg/d encorafenib (**E-G**). Cardiac function and dimensions were measured by echocardiography. **A**,**E**, M-mode images from short-axis views of the heart were analysed to provide data for cardiac dimensions. Representative images are in the upper panels with analysis of echocardiograms shown below. LV, left ventricle; AW, anterior wall; PW, posterior wall; ID, internal diameter. Diastolic measurements are shown with all data provided in **Supplementary Table S10. B**,**F**, B mode images from mice at 3 d (upper panels) or 7 d (lower panels) were subject to strain analysis using speckle-tracking to gain an estimate of left ventricular mass. EDLVM, end diastolic left ventricular mass. Additional data are provided in **Supplementary Table S9. C**,**G**, Haemotoxylin (H&E) and eosin staining of mouse heart sections. Representative images are on the left with quantification of cardiomyocyte size from H&E-stained sections on the right. Individual data points are shown with means ± SEM. Statistical analysis used a two-tailed paired t test. **D** and **H**, RNA was isolated from hearts of mice treated with vehicle, SB590885 or encorafenib for 7 d, and mRNA expression assessed by qPCR. Data are individual points with means ± SEM.

We next assessed the effects of encorafenib on mouse hearts *in vivo*. Encorafenib is used at 300-450 mg/d in humans, so we selected a dose of 3 mg/kg/d for delivery by osmotic minipumps. Like SB590885, encorafenib increased systolic/diastolic anterior and posterior wall thickness (**Figure 6E; Supplementary Table S10**) and left ventricular mass (**Figure 6F**). Encorafenib also increased cardiomyocyte cross-sectional area (**Figure 6G**), consistent with cardiomyocyte hypertrophy without any increase fibrosis, and it did not affect expression of *Nppa* or *Nppb* mRNAs (**Figure 6H**). We conclude that Type 1 RAF kinase inhibitors promote cardiomyocyte hypertrophy *in vivo*, consistent with their effects in cultured cardiomyocytes. *In vivo*, the drugs increase cardiomyocyte cross-sectional area, in the absence of pathological features of cardiac hypertrophy such as fibrosis or inflammation.

## Discussion

BRAF signalling has been studied extensively in relation to cancer because of its oncogenic potential, but the role of BRAF in the heart has remained relatively unexplored. As illustrated in **Figure 7**, this study showed that BRAF is upregulated in human heart failure and activation of BRAF in cardiomyocytes promotes cardiomyocyte hypertrophy. The data for BRAF in adult human heart failure (**Figure 1**) are intriguing, although (as with any studies of end-stage disease) it is difficult to know whether the changes are cause or consequence of the disease. Upregulation of BRAF in separate cohorts of heart failure patients, with selective upregulation of *BRAF* and downregulation of *RAF1* and *ARAF* isoforms in patients with dilated cardiomyopathy, suggests that the changes are a key feature of a failing heart. Even if BRAF is not a driving force in the development of human heart failure, an increase in BRAF expression (and potentially, BRAF activation) may influence the failing phenotype. It should be noted that RAF kinases are potentially expressed at different ratios and levels in different cell types, so the differences in expression of the RAF isoforms could simply result from changes in cellular content in failing hearts compared with the normal controls. In this case, the changes may sustain a failing phenotype and reduce potential for recovery. On the other hand, an increase in BRAF expression may reflect an increase in cardioprotective signalling. On balance, it seems appropriate to consider that the increase in BRAF expression in heart failure is potentially an important facet of the disease.

**Figure 7.**
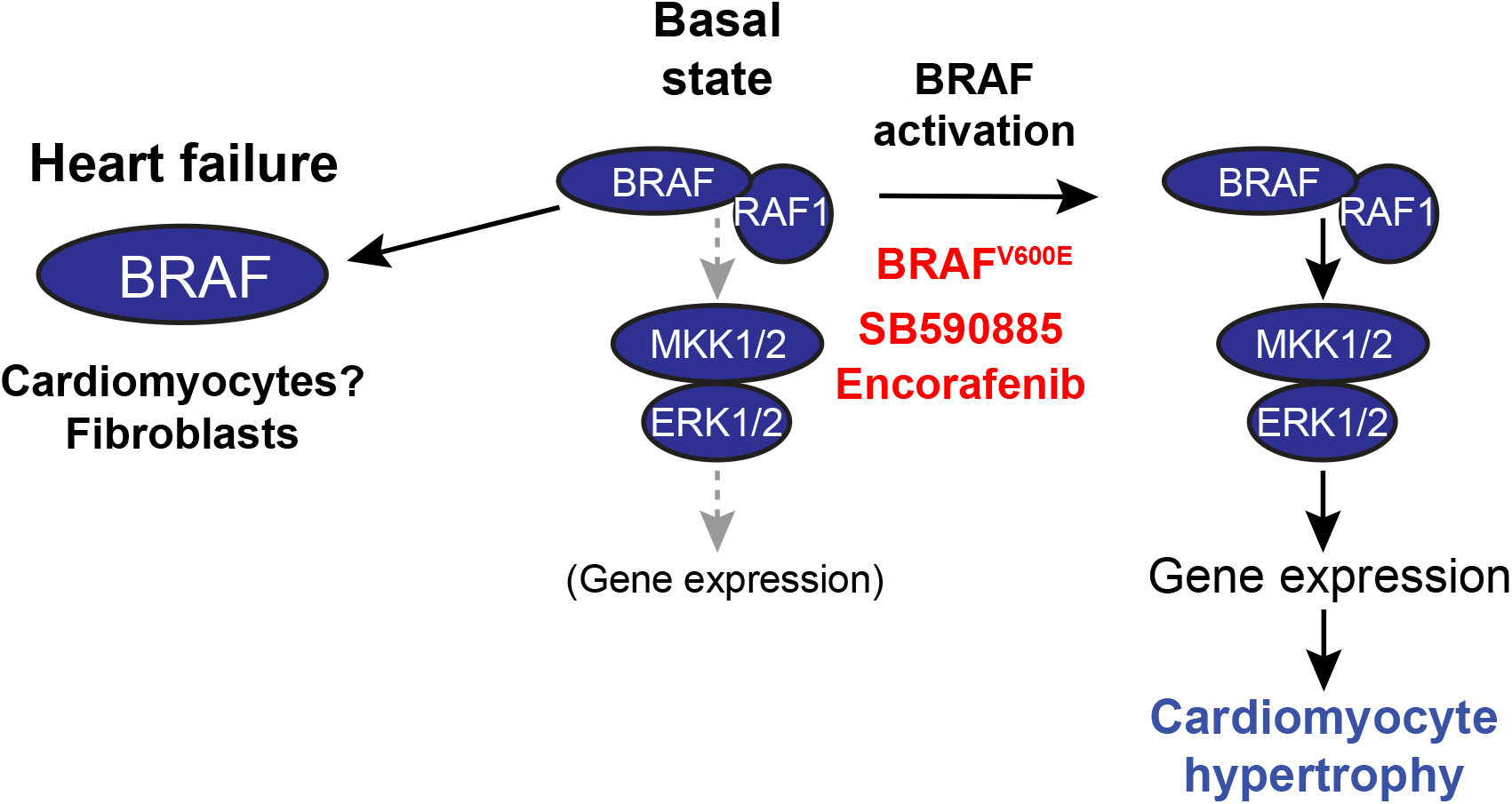
Schematic representation of the conclusions from this study. BRAF is upregulated in human failing hearts, although the cells involved remain to be defined. In cardiomyocytes, BRAF and RAF1 exist in preformed complexes. Activation of cardiomyocyte BRAF by knock-in of the V600E mutation or with Type 1 RAF kinase “inhibitors” (e.g. SB590885, encorafenib) results in activation of MKK1/2 (mitogen-activated protein kinase kinases 1 and 2) and ERK1/2 (extracellular signal-regulated kinases 1 and 2) with stimulation of downstream gene expression resulting in cardiomyocyte hypertrophy.

ERK1/2 were shown to be activated in cardiomyocytes by hypertrophic agonists as early as 1993 [39]. Subsequent work from many groups established that ERK1/2 signalling is important in cardiomyocyte hypertrophy, but also that it is cardioprotective [6-8]. Possibly the earliest publication on RAF kinases in the heart was in 1995 [10], but this focused on RAF1 and ARAF, the difficulty with BRAF being the lack of antibodies of sufficient quality to study BRAF properly in cardiac tissues. Although this was discussed, it became generally assumed that BRAF was not expressed at any significant levels in the heart. Assessment of the cardiomyocyte kinome showed quite clearly that BRAF is, indeed, expressed in cardiomyocytes [15]. Here, we show further that activation of cardiomyocyte BRAF alone is capable of driving hypertrophy, since activation of endogenous BRAF by knock-in of the V600E oncogenic mutation promotes cardiac and cardiomyocyte hypertrophy *in vivo* (**Figures 2** and **3**). Although there was increased mRNA expression of foetal gene markers (*Myh7, Nppa, Nppb*) and pro-fibrotic genes (*IL11, Col1a1*) generally associated with pathological hypertrophy, histological assessment of the myocardium showed little indication of any fibrosis (**Figure 3**). Indeed, the increase in left ventricular wall thickness and predicted mass appeared to be attributable solely to cardiomyocyte hypertrophy, with increased cardiomyocyte cross-sectional area and an associated enhancement of fractional shortening and ejection fraction. These data are consistent with compensated, concentric hypertrophy in mice with cardiomyocyte-specific expression of constitutively-active MKK1 [40], and in accord with studies of mice with deletion of ERK1 and/or ERK2 in heart failure models showing that ERK1/2 signalling is cardioprotective and required for concentric hypertrophy [41].

Activating the ERK1/2 cascade in cardiomyocytes may be beneficial in pathological situations that lead to heart failure. However, activating the pathway in fibroblasts is potentially less desirable since it may increase cardiac fibrosis [34, 42]. Harnessing the benefits of the ERK1/2 pathway for the heart is therefore challenging: activation of protein kinase signalling is less straightforward than inhibition, and selective activation in one cell type over another is even more difficult. Our data suggest that Type 1 RAF inhibitors may be a way to do this. Thus, the established Type 1 inhibitor SB590885 activated ERK1/2 signalling in cultured cardiomyocytes and perfused hearts (**Figure 4G-J**), increasing cardiomyocyte cross-sectional area in cells in culture (**Figure 5D)** and in hearts *in vivo* (**Figure 6C**). Similar effects were seen with encorafenib, a BRAF-targeted inhibitor used clinically, albeit generally in combination with a MKK1/2 inhibitor [38]. Our data suggest encorafenib is a Type 1 inhibitor, since it activated ERK1/2 (**Figure 5E**) and increased cardiomyocyte size both in cultured cells (**Figure 5D**) and *in vivo* (**Figure 6G**). The effects of SB590885 and encorafenib contrast with dabrafenib, a Type 1.5 inhibitor that, in our hands, has limited RAF paradox-inducing effects and does not induce cardiomyocyte hypertrophy in wild-type mice [21]. Indeed, dabrafenib inhibited fibrosis in the hearts of mice treated with angiotensin II to induce hypertension, suggesting that it serves as an inhibitor of signalling in the heart, rather than an activator. The mechanisms behind “RAF paradox” signalling in cancer cells are reasonably well-investigated [20]: Type 1 inhibitors bind to RAF kinases in an active conformation, increasing dimerisation potential; Type 1.5 inhibitors bind a similar conformation, but with some alterations more consistent with an inactive enzyme. This could explain why SB590885/encorafenib and dabrafenib have different effects on the heart.

As terminally-differentiated cells, cardiomyocytes may be viewed as having “hard-wired” ERK1/2 signalling. This is reflected in the pre-formed RAF complexes detected in cardiomyocytes even from neonatal hearts (**Figure 4A-E**), suggesting that cardiomyocytes are primed for RAF signalling, enabling them to respond rapidly to extracellular stimuli. Consistent with this, the ERK1/2 cascade can be activated within 3 min of stimulation with receptor agonists (e.g. epidermal growth factor or endothelin-1 [11]), with upregulation of the first ERK1/2-dependent immediate early genes within 15-30 min [43]. Such rapid signalling may be facilitated by pre-formed signalling complexes. Such “signalosomes” are established for the ERK1/2 cascade itself [44], and the presence of ERK1/2 signalosomes in cardiomyocytes could account for rapid activation in association with receptor stimulation, but also allow for the relatively slow effect of SB590885 which induces BRAF paradox signalling over 10-20 min (**Figure 4H-I**). For the latter, the drug needs to diffuse into cells and permeate intracellular compartments before becoming effective. SB590885 not only activated ERK1/2 in the cytoplasm, but also increased activated ERK1/2 in the nucleus (**Figure 5A,B**). As in our previous studies of endothelin-1 [33], we did not detect net translocation of ERK1/2 to the nucleus following stimulation with SB590885. Interestingly, we had greater relative activation in the nucleus than the cytoplasm compared with endothelin-1, potentially resulting from free diffusion of the drug rather than receptor-driven signalling. This raises the intriguing possibility that the ERK1/2 cascade in cardiomyocytes is hard-wired to such a degree that ERK1/2 signalling is strictly compartmentalised, and signalling in the nucleus is divorced from that in the cytoplasm. Consistent with the presence of nuclear-activated ERK1/2, SB590885 promoted ERK1/2-dependent changes in immediate early gene expression and induced an increase in cardiomyocyte size (**Figure 5C,D**). This is in accord with the body of evidence implicating ERK1/2 signalling in cardiomyocyte hypertrophy [45].

Anti-cancer therapies are designed to prevent cell growth/proliferation and promote death of cancerous cells, and this has driven much of the pioneering research into intracellular signalling and regulation of cell function. The requirements for cancer therapies are generally diametrically opposed to the requirements of an adult heart subjected to pathophysiological stresses, in which cytoprotection and increased cardiomyocyte growth (if not proliferation) are desirable [46]. It is therefore unsurprising that cancer therapies have cardiotoxic effects both during cancer treatment and in cancer survivors [47, 48]. MKK1/2 inhibitors clearly have cardiotoxic effects and trametinib causes hypertension in ∼26% of patients with decreased ejection fraction in 7-11% of patients [49, 50]. This is likely to be an on-target effect since alternative MKK1/2 inhibitors (selumetinib and cobimetinib) are also associated with increased risk of hypertension and reduced ejection fraction [51]. The situation with RAF inhibitors is less clear. Vemurafenib increases QT interval [42], but dabrafenib monotherapy has not been associated with any obvious cardioxicity [52, 53], suggesting that the effects of vemurafenib are unrelated to inhibition of RAF kinases. Encorafenib has not yet been used extensively as a monotherapy, being used clinically in combination with the MKK1/2 inhibitor binimetinib [38], but our data suggest that, by activating the “RAF paradox” in cardiomyocytes, encorafenib is more likely to be beneficial for the heart, rather than detrimental. With any small molecule inhibitor, there is always consideration of off-target effects. However, we used two different compounds to activate the “RAF paradox” (SB590885 and encorafenib), both of which promoted cardiomyocyte hypertrophy *in vitro* and *in vivo* (**Figures 4-6**), suggesting the response is an on-target effect.

Overall, this study provides data to demonstrate that selective activation of ERK1/2 signalling in cardiomyocytes (potentially desirable in a failing heart) can be achieved. The Type 1 RAF inhibitors being used for cancer are not cardiotoxic *per se* and, with further research, they may even prove useful as a supporting therapy in failing hearts. In this respect, it is necessary to consider whether using drugs to enhance ERK1/2 signalling for heart failure could increase the potential for unregulated cell proliferation or even cancer. One side-effect of RAF inhibitors is the development of squamous cell carcinoma [16], but this is manageable in the context of cancer therapy. It may also be appropriate to limit the duration of treatment to restrict the possibility of cancers developing. However, heart failure is just as devastating a disease as cancer, and the benefits afforded potentially outweigh the risks in this setting.

## Data Availability Statement

All primary data are available from the corresponding author upon reasonable request. Additional data sharing information is not applicable to this study.

## Abbreviations

ERK1/2: extracellular signal-regulated kinase
MAPK: mitogen-activated protein kinase
MKK: MAPK kinase

