## Supplemental Tables for "Cardiomyocyte BRAF and type 1 RAF inhibitors promote cardiomyocyte and cardiac hypertrophy in mice *in vivo*"

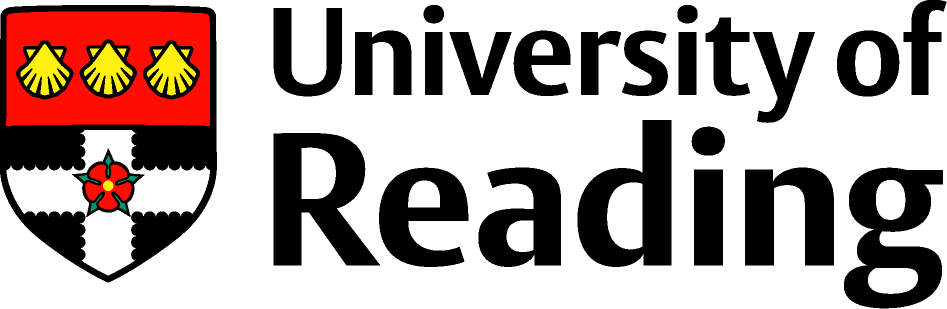

Professor Mark Lemmon

Chair of the Editorial Board,

Biochemical Journal

10^th^ August 2021

Dear Professor Lemmon,

We wish to submit our manuscript “Cardiomyocyte BRAF and type 1 RAF inhibitors promote cardiomyocyte and cardiac hypertrophy in mice *in vivo*” to *Biochemical Journal* for consideration for publication as a Research Paper.

In this manuscript, we show that BRAF is upregulated in heart samples from failing human hearts compared with normal control hearts. We then demonstrate that activation of endogenous BRAF in mouse cardiomyocytes by knock-in of the oncogenic V600E mutation promotes cardiomyocyte hypertrophy *in vivo*. Thus, cardiomyocyte BRAF is sufficient to induce hypertrophy. Finally, we explored the effects of Type 1 RAF inhibitors on ERK1/2 signalling in cultured cardiomyocytes and in mouse hearts *in vivo*, demonstrating that cardiomyocytes are prone to activation of ERK1/2 by such compounds via the “RAF paradox”, and these compounds lead to cardiomyocyte and cardiac hypertrophy. At a biochemical level, the study has implications for understanding of RAF signalling in the heart. There are translational implications since this may provide a mechanism for boosting cardiac function as hearts begin to fail. In relation to the cardio-oncology field, the data demonstrate that at least one of the licensed anti-cancer RAF inhibitors in clinical use (encorafenib) activates the RAF paradox in the heart, promoting hypertrophy. Moreover, this drug alone appears to have no overt cardiotoxic effects.

I look forward to hearing from you.

Yours sincerely,

Angela Clerk
